# Visual Occlusions Result in Phase Synchrony Within Multiple Brain Regions Involved in Sensory Processing and Balance Control

**DOI:** 10.1101/2022.10.28.514058

**Authors:** Evangelia-Regkina Symeonidou, Daniel P. Ferris

## Abstract

There is a need to develop appropriate balance training interventions to minimize the risk of falls. Recently, we found that intermittent visual occlusions can substantially improve the effectiveness and retention of balance beam walking practice (Symeonidou and Ferris 2022). We sought to determine how the intermittent visual occlusions affect electrocortical activity during beam walking. We hypothesized that areas involved in sensorimotor processing and balance control would demonstrate spectral power changes and inter-trial coherence modulations after loss and restoration of vision. Ten healthy young adults practiced walking on a treadmill-mounted balance beam while wearing high-density EEG and experiencing reoccurring visual occlusions. Results revealed spectral power fluctuations and inter-trial coherence changes in the visual, occipital, temporal, and sensorimotor cortex as well as the posterior parietal cortex and the anterior cingulate. We observed a prolonged alpha increase in the occipital, temporal, sensorimotor, and posterior parietal cortex after the occlusion onset. In contrast, the anterior cingulate showed a strong alpha and theta increase after the occlusion offset. We observed transient phase synchrony in the alpha, theta, and beta bands within the sensory, posterior parietal, and anterior cingulate cortices immediately after occlusion onset and offset. Our results provide support for cross-modal phase resetting and enhanced processing in areas involved in sensory processing and balance control as an explanation for increased long-term balance improvement when training with intermittent visual occlusions. Our training intervention could be implemented in senior and rehabilitation centers, improving the quality of life of elderly and neurologically impaired individuals.

## Introduction

Visual perturbations can enhance balance training efficacy in a beam walking task and decrease the risk of falls (Peterson et al., 2018; Peterson & Ferris, 2018; Symeonidou & Ferris, 2022). A previous study in our lab showed that healthy, young participants practicing beam walking with brief reoccurring visual rotations had a higher balance improvement compared to the group without visual rotations (Peterson et al., 2018). The group training with intermittent visual rotations had an immediate improvement in step-offs off the beam of 42% compared to the group without visual rotations’ improvement of 9%. In a more recent study (Symeonidou & Ferris, 2022), we used intermittent visual occlusions as a beam walking training intervention. Participants who practiced with the visual occlusions showed long-term balance improvements compared to the control group. On the same day of the training the visual occlusions group had a 78% improvement in step-offs, an almost four-fold difference compared to the participants training without occlusions who experienced only a 21% reduction in step-offs. At a retention testing of the same task two-weeks later, the visual occlusions group still had a 60% improvement in step-offs compared to the 5% improvement of the control group. These results suggest that visual occlusions can induce long-term balance improvements in a beam-walking task.

To better integrate visual occlusions in balance training interventions we need to understand the underlying changes in cortical activity when the perturbations are presented. The electroencephalography (EEG) data in Peterson and Ferris (2018), indicated that visual rotations were accompanied by theta (4-8 Hz), alpha (8-13 Hz), and beta (13-30 Hz) spectral power changes in parietal and occipital cortices. These spectral modulations provided evidence of the occipital and posterior parietal cortex involvement in the response to the visual perturbations during dynamic balance control.

The posterior parietal cortex plays a key role in integrating and storing visual information required for step planning (Drew & Marigold, 2015; Peterka, 2002). In an obstacle stepping task, cats showed increased posterior parietal cortex firing when approaching the object, they would step over (Lajoie et al., 2010). Some posterior parietal cortex cells continued firing even when vision was occluded during that time period (Marigold & Drew, 2011). Lesions within this cortical area led to working memory decrease of the object’s properties and disrupted step planning by incorrect height adjustment of the hind legs when stepping over it (McVea et al., 2009). The above data suggests that the posterior parietal cortex can store visual and proprioceptive information for step planning during locomotion when visual information is not available.

We can determine if visual occlusions during beam walking increase communication and processing within the sensory and posterior parietal cortices by estimating phase modulations within these areas. Visual and other stimuli can induce phase synchrony within and across cortical regions (Kayser et al., 2008; Lakatos et al., 2008; Rajkai et al., 2008; Schroeder & Lakatos, 2009; Voloh et al., 2015). This enhances communication between neural populations, so that incoming information reaches the target site during a high excitability phase, improving processing, neuroplasticity, and learning (Fries, 2005; Nash-Kille & Sharma, 2014; Womelsdorf et al., 2007). In multisensory events, sensory cues of one modality synchronize and reset the oscillatory phase to improve processing of upcoming events of another sensory modality (Kayser et al., 2008; Lakatos et al., 2008; Thorne et al., 2011). This is not specific to multisensory cues; oscillatory phases can also be modulated by sensory, motor, and attentional cues to affect processing of upcoming events (Lakatos et al., 2013; Makeig et al., 2004; Rajkai et al., 2008; Shah et al., 2004; Van Atteveldt et al., 2014). To determine phase synchrony, we will need to compute the EEG trial-to-trial phase relationship by Inter-trial coherence (ITC) estimation. Estimating ITC and spectral power changes relative to the occlusion event can inform us how visual occlusions affect the amplitude and phase of neural oscillatory activity within and across cortical regions relevant to dynamic balance control.

The goal of this study was to determine if intermittent visual occlusions induce increased cortical phase synchrony doing practice of a whole body motor task (Fell & Axmacher, 2011). We had ten healthy young participants practice walking on a treadmill-mounted balance beam while liquid crystal glasses provided transient, intermittent visual occlusions. We recorded high-density scalp EEG to investigate changes in cortical processing related to the intermittent visual occlusions. We hypothesized that visual occlusions would induce spectral power changes and phase synchrony in areas relevant to sensory processing, motor planning and multisensory integration. Specifically, we expected changes within the occipital, temporal, sensorimotor, and posterior parietal cortex, reflecting increased sensory processing for subsequent step planning when vision is absent, leading to balance improvement during beam walking. We based our hypothesis on previous EEG data by Peterson and Ferris (Peterson & Ferris, 2018) and the additional presumption that an increase in phase synchrony would improve processing of sensory feedback relevant to successfully maintaining balance (Fell & Axmacher, 2011; Lakatos et al., 2008; Rajkai et al., 2008; Thorne et al., 2011).

## Materials and Methods

Ten young, healthy, and right leg dominant participants (6 males, 4 females, age = 24.3 ± 5.1) took part in this study. We assessed leg dominance by asking participants which foot they would kick a ball with. Participants had no neurological, orthopedic, and musculoskeletal conditions, or lower limb surgeries. The study was approved by the University of Florida Institutional Review Board (IRB) and was conducted according to the Declaration of Helsinki. All participants had to provide written informed consent prior to their participation in this study.

Participants practiced tandem walking on a treadmill mounted balance beam while wearing liquid crystal lens glasses (Senaptec Strobe, Senaptec, Oregon, USA) and experiencing intermittent visual occlusions as described in Symeonidou and Ferris (2022) (Fig. 1). Briefly, the experiment consisted of three 10-minute training sessions (30 minutes total) and a 3-minute pre and post-test session, prior and after the training. The balance beam was 2,5 cm high and 2,5 cm wide and participants walked at a fixed speed of 0.22 m/s. The visual occlusions were presented during the training session in a reoccurring fashion as follows: 1.5 s of occlusion followed by 7.5 s of clear vision. For occlusion presentation, we used the Senaptec app (Senaptec, Oregon, USA) on a Samsung Note 10 (Samsung, Seoul, South Korea) and the occlusion cycle was initiated and controlled by a custom MATLAB script. We initiated the cycle every time participants stepped on the beam and paused it when they stepped off the beam. For the occlusion cycle, we used a variable delay up to 1 s to avoid any cognitive anticipatory effects.

**Figure 1.**
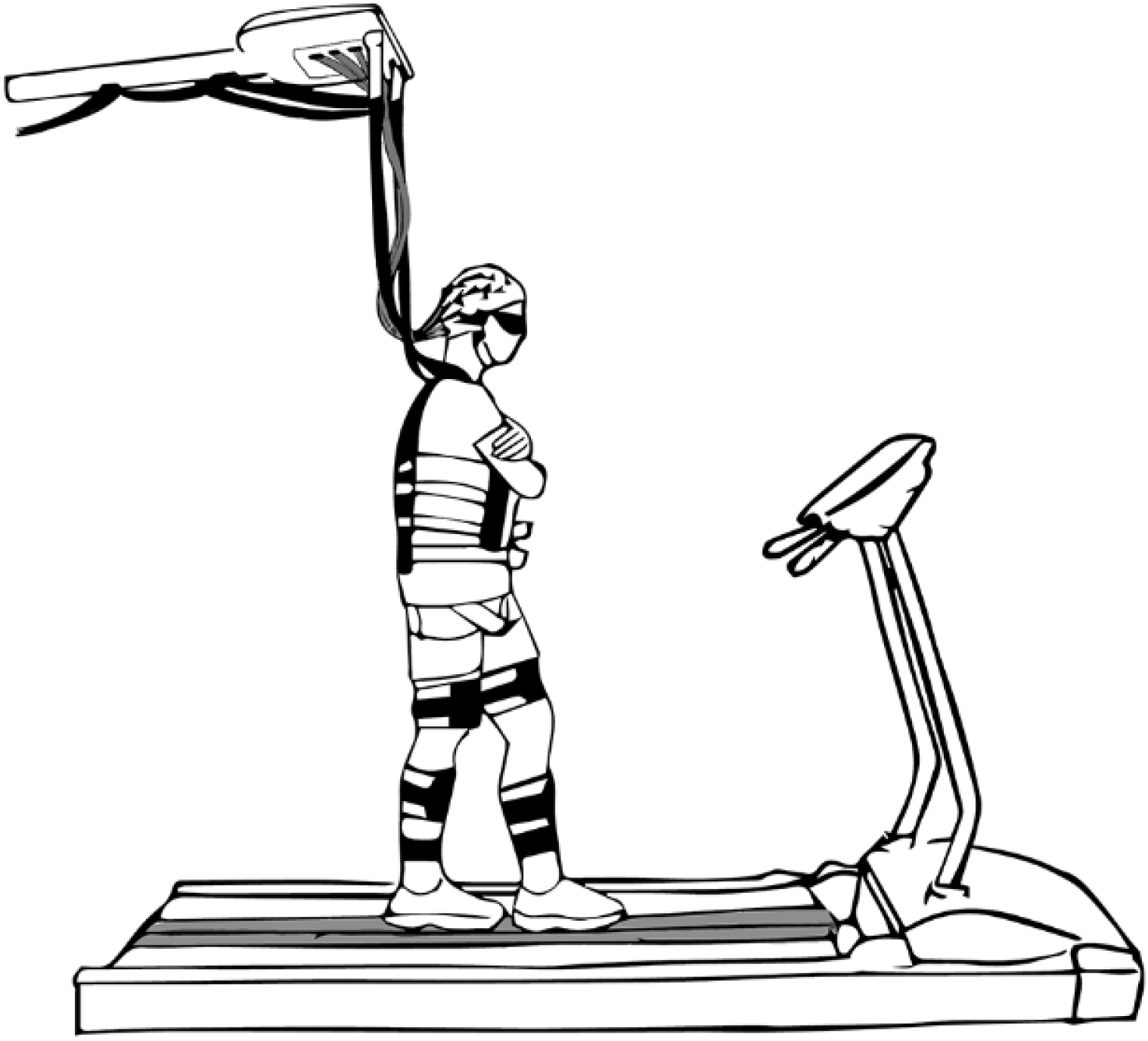
Experimental Setup. A sketch of a participant walking on the treadmill-mounted balance beam while wearing the occlusion glasses and high-density EEG. The EEG analog to digital converter and battery were placed on top of a body weight support system. To minimize electrode cable movement, cables were bundled together and suspended from the body weight support system. Participants wore a harness for safety that allowed for freedom of movement in the mediolateral direction.

We recorded EEG to track neurophysiological changes relative to the visual occlusions. We used a 128-channel BioSemi ActiTwo system (BioSemi, Amsterdam, The Netherlands) to record electrical scalp activity. We also placed 8 external electrodes on the participants’ neck muscles -one on the splenius capitis, one on the levator scapulae, and two on the trapezius of each side to record electrical neck muscle activity and sampled the data at 512 Hz. To reduce motion artifacts caused by excessive cable sway (Symeonidou et al., 2018), we placed the EEG amplifier on top of the body-weight support system and wrapped the suspended EEG cables together. We adjusted the height of the body-weight support system for each participant and made sure there is no cable pulling when participants swayed in the mediolateral direction. To avoid pulling in the anteroposterior direction participants were instructed to stay within a designated area marked on the treadmill. To determine subject-specific electrode locations, we used the Artec Eva structured light 3d scanner (Artec3D Santa Clara, CA, USA) and created a 3D model of the participants head and electrodes (Shirazi & Huang, 2019).

To analyze changes in brain activity around the occlusion event, we recorded the timing of the occlusion onset and offset with a light sensor. We used a 5×3×6.4 mmBPW46 PIN photodiode (Vishay Electronic, Selb, Germany) that we attached to the inner side of the outer corner of the right lens. To increase the voltage change occurring when the glasses changed transparency, the photodiode was connected to a circuit with an LM324 operational amplifier (Dallas, Texas, US). We recorded a voltage change from 1 V to 1.8 V when glasses changed from opaque to transparent. The amplifier circuit was attached to the participant’s harness and was connected to a NI-DAQxm (National Instruments, Austin Texas). To sync the photodiode signal with the EEG recording we used a 2 Hz square wave that was also connected to the NI-DAQxm. Both signals were recorded with Motive 2.1 (Natural Point Inc, Corvallis, Oregon, USA) at 1000 Hz.

Events from the occlusion onset and offset were imported into the EEG training sessions using timestamps of the square wave rising edge to align the signals. The training sessions were then concatenated into one data file. The data were down sampled to 256 Hz and electrode locations were added using the get_chanlocs toolbox in EEGLAB (v14.1.2). We subsequently used a high-pass filter at 1 Hz and re-referenced the signal to a common median reference in order to increase the signal to noise ratio for the subsequent steps of the analysis. We used the clean line extension within EEGLAB to remove any line noise and removed channels if they: a) had a standard deviation (SD) > 1000 μV, b) had kurtosis >5 SDs or c) were uncorrelated > 1 % of the time with the other channels. We also removed channels based on manual inspection. We performed artifact subspace reconstruction (ASR) with a SD of 20 using a 5 min pre-recorded EEG baseline where the participant was standing still on the treadmill with the glasses on the transparent modus (Mullen et al., 2013; Nordin et al., 2020; Peterson & Ferris, 2018). This baseline was pre-processed in the exact same way we processed the training dataset. In a subsequent step, we used empirical mode decomposition (EMD) to find the 1st intrinsic mode function (IMF) of the signal and then applied canonical correlation analysis to remove the components whose variance exceeded the median interquartile range of that IMF. The 1st IMF contains the high-frequency portion of the signal, which is suggested to be muscle or other type of artifact and not brain-related activity (Peterson & Ferris, 2018). Next, we interpolated the missing electrodes and average referenced the EEG and EMG electrodes separately. After reducing the data to 80 principal components, we performed independent component analysis (ICA) using the AMICA algorithm, which was also set to reject any bad frames.

After acquiring the weights of the independent components, we epoched the EEG data around the perturbation onset (-500 ms, +2000 ms) and performed dipole fitting using the DIPFIT function within EEGLAB (Oostenveld & Oostendorp, 2002). We modelled each independent component as an equivalent current dipole within a boundary element head model using subject specific magnetic resonance images for the participants of the occlusions group. Two participants were removed from the analysis as they had missing EEG data from parts of the training session (due to cables getting disconnected or the battery dying in the middle of the recording). For all remaining 8 participants, we retained components that explained >85% of the scalp variance and clustered the dipoles using k means clustering with the following weighting: 2 for spectral power, 10 for dipole locations, and 1 for scalp maps. Dipoles that were >3 SD away from the rest of the clusters were marked as outliers and discarded. We also discarded clusters that contained dipoles from less than five participants and only included one dipole per participant per cluster. The selection of the dipole for each participant was based on manual inspection of their scalp map, spectral power, and distance from the cluster centroid. We retained clusters in the right superior occipital, left superior occipital, right inferior occipital, left inferior occipital, posterior parietal, temporal, sensorimotor, posterior cingulate, anterior cingulate, and prefrontal cortex, and the inferior frontal gyrus. We plotted event-related spectral perturbation (ERSP) graphs using the median across trials within each participant and by averaging across participants and within cluster. The ERSPs were normalized using a pre-stimulus baseline of -500 ms to 0 ms relative to the visual occlusion onset and significance masked using bootstrap statistics. We also plotted the inter-trial coherence (ITC) plots for the above clusters, to identify if neural phase resetting occurred within-cluster after the occlusion presentation. The ITC values were calculated similarly to the ERSP values. They were normalized using a pre-stimulus baseline of -500 ms to 0 ms.

## Results

EEG analysis revealed eleven brain areas with broad electrocortical activity related to the intermittent visual occlusions during beam walking (Fig 2). Our event-related spectral perturbation (ERSP) plots showed alpha synchronization in the occipital, posterior parietal, temporal, sensorimotor, and posterior cingulate cortex ∼500 ms after the occlusion onset (Fig 3). The right superior occipital and posterior parietal cortex exhibited theta (6-8 Hz), alpha (8-13 Hz) and beta (13-30 Hz) synchronization immediately after vision was restored. The posterior parietal cortex also showed theta desynchronization during most of the occlusion duration. The temporal cortex exhibited theta synchronization after the occlusion onset and offset, as well as beta synchronization ∼1000 ms into the occlusion. The anterior cingulate showed a similar theta pattern, with high synchronization after the occlusion onset and offset. The prefrontal cortex showed gamma desynchronization (30-100 Hz) during the occlusion that persisted up to ∼500 ms after the occlusion offset.

**Figure 2.**
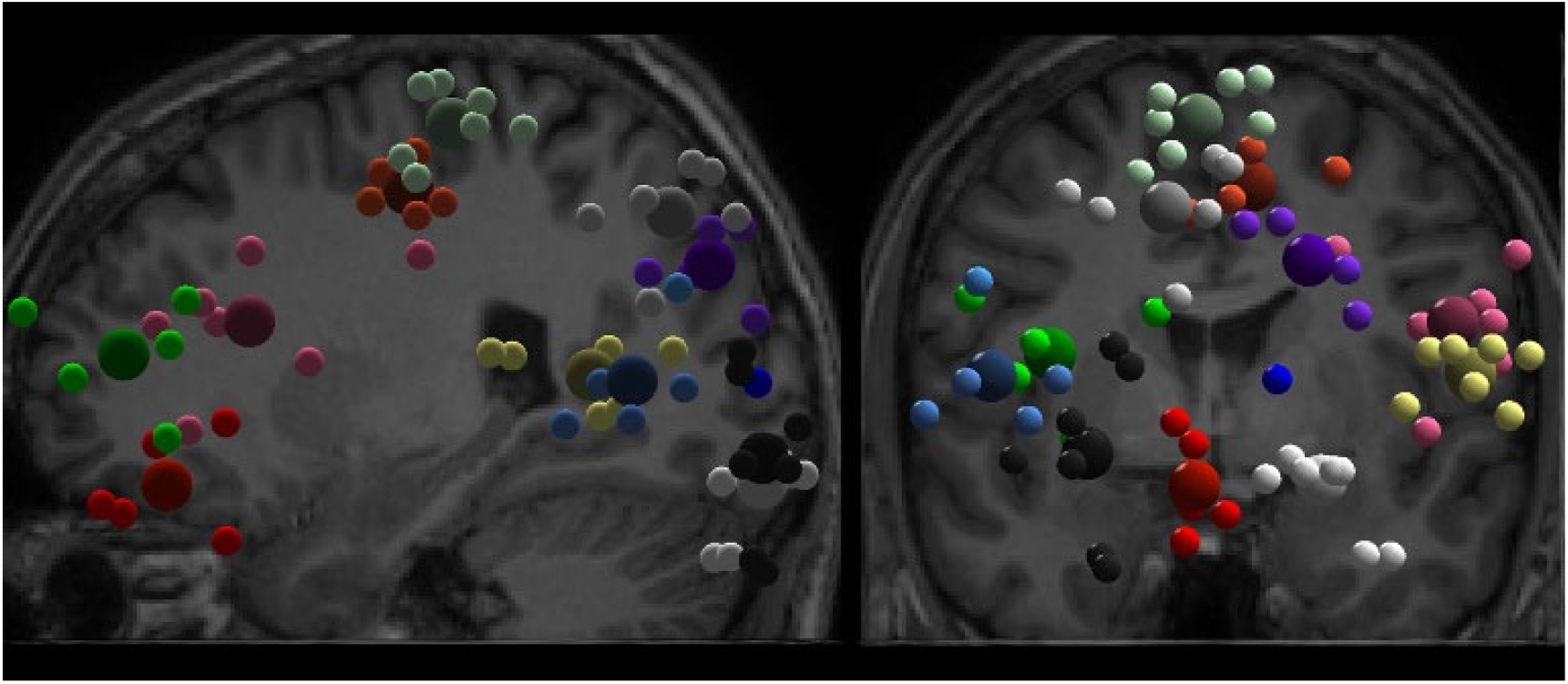
Brain sources. Brain sources exhibiting activity changes during beam walking practice with intermittent visual occlusions (n=8). Cortical clusters containing cluster centroids (big spheres) and participant dipoles (small spheres) in sagittal (left) and coronal (right) plane. Coloring: the prefrontal cortex is depicted in green, the anterior cingulate in red, the inferior frontal gyrus in magenta, the posterior cingulate in orange, the sensorimotor cortex in mint, the temporal cortex in yellow, the posterior parietal cortex in gray, the right superior occipital cortex in purple, the left superior occipital cortex in blue, the left inferior occipital cortex in black, the right inferior occipital cortex in white.

**Figure 3.**
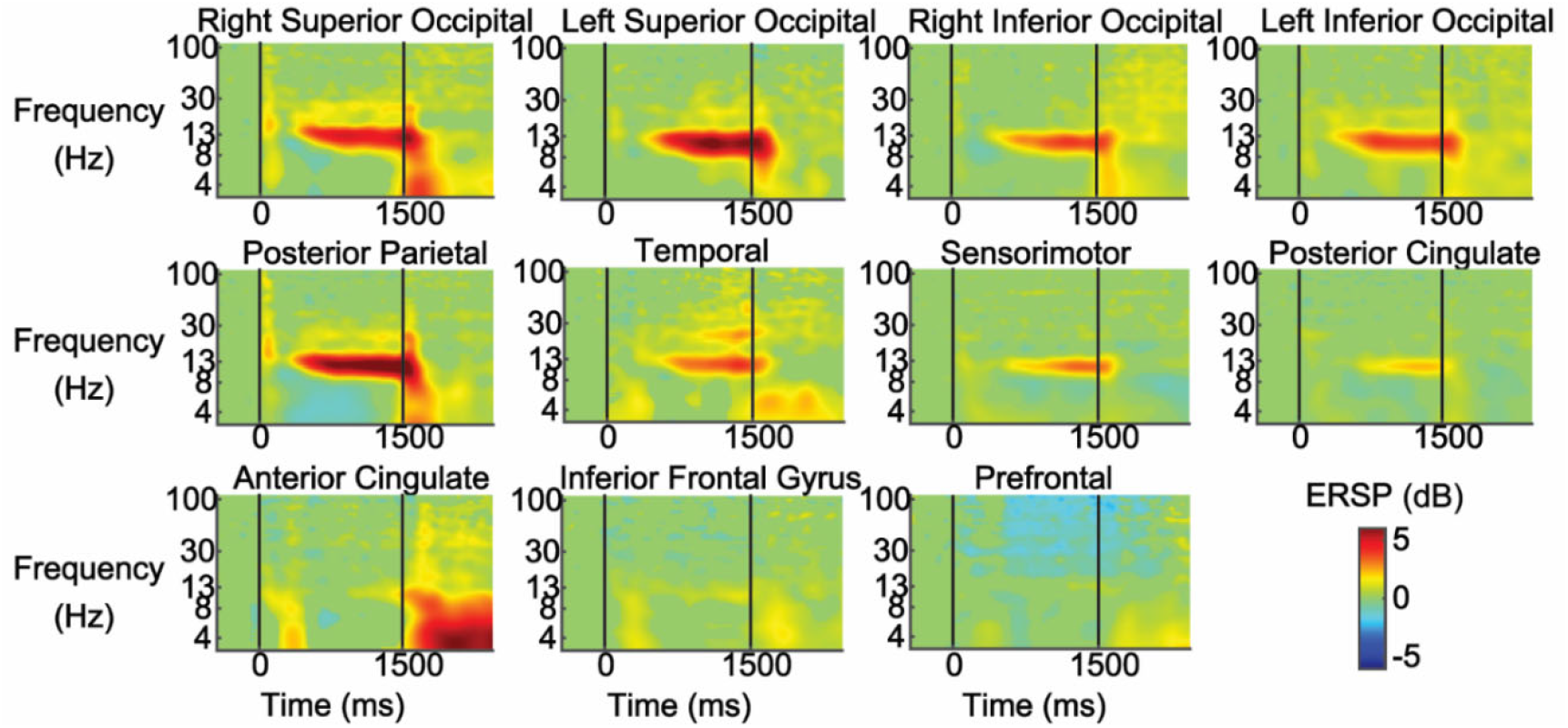
Event-related spectral perturbation plots. Event-related spectral perturbation plots (ERSPs) epoched around the visual occlusion onset (0 ms) and offset (1500 ms). The ERSPs are shown for the right and left superior occipital cortex, right and left inferior occipital cortex, posterior parietal cortex, temporal cortex, sensorimotor cortex, posterior cingulate cortex, anterior cingulate cortex, inferior frontal gyrus, and prefrontal cortex. Red reflects synchronization and blue reflects desynchronization. Non-significant ERSP power changes were set to 0 dB (green). Visual Occlusion resulted in widespread alpha synchronization in occipital, parietal, and temporal brain regions. The pre-frontal cortex showed gamma desynchronization during the occlusion and the anterior cingulate showed strong theta synchronization after restoration of vision.

The inter-trial coherence plots (ITC) (Fig 4) revealed an increased coherence with visual occlusion in a wide number of clusters. In the occipital clusters, we saw increased coherence in the theta and alpha frequencies after the occlusion onset and in the theta frequency after the occlusion offset. The effect was the strongest in the right superior occipital cluster. A similar pattern was observed in the anterior cingulate. The posterior parietal cortex showed a prominent beta coherence increase after occlusion onset and a theta and alpha increase after vision was restored. The temporal, sensorimotor cortex, and inferior frontal gyrus showed an alpha and low beta coherence increase after the occlusion onset, and a less pronounced theta coherence increase after the occlusion offset. Last, there was an increase in theta and alpha coherence after occlusion onset and offset in the posterior cingulate.

**Figure 4.**
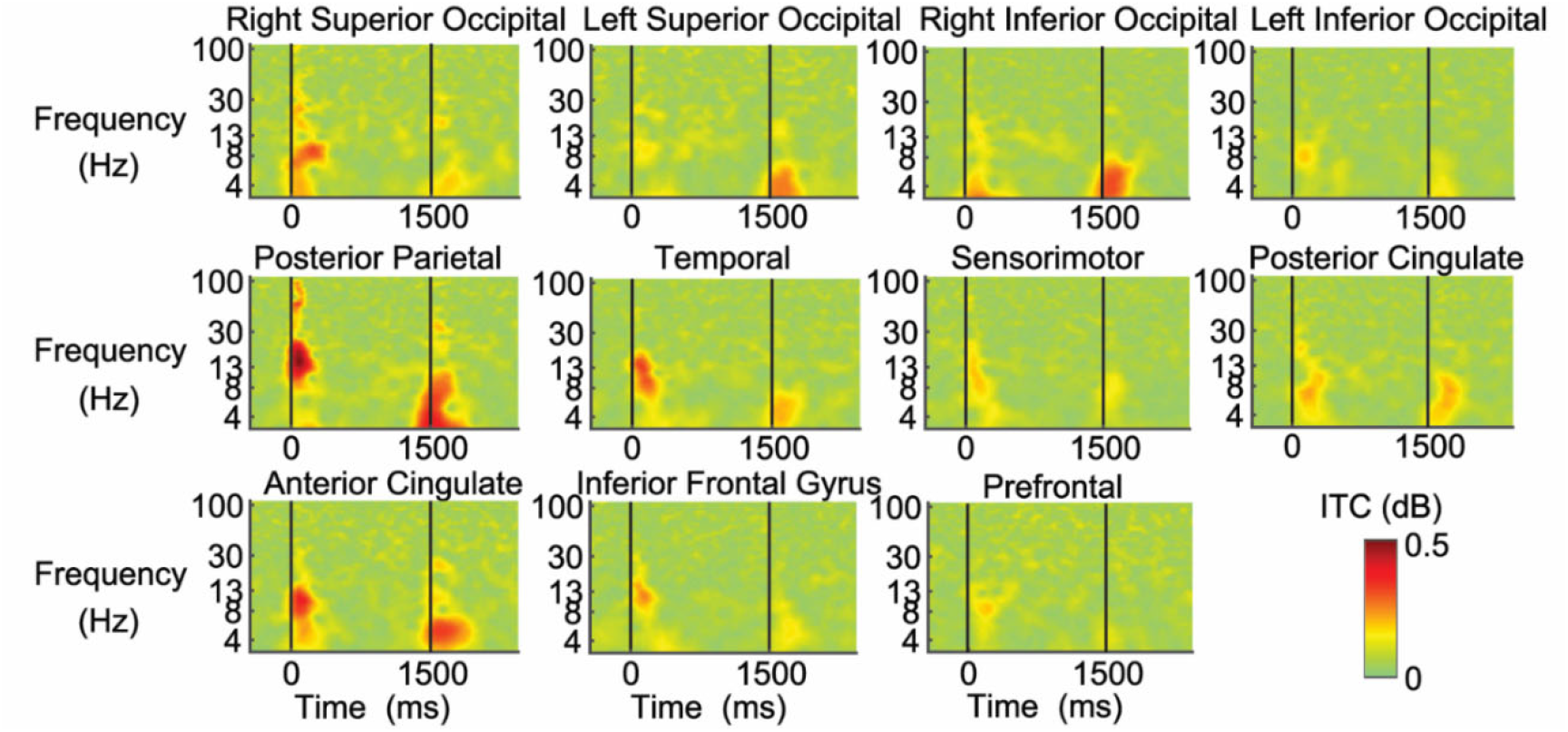
Inter-trial coherence plots. Inter-trial coherence (ITC) plots epoched around the visual occlusion onset (0 ms) and offset (1500 ms). The ITCs are shown for the right and left superior occipital cortex, right and left inferior occipital cortex, posterior parietal cortex, temporal cortex, sensorimotor cortex, posterior cingulate cortex, anterior cingulate cortex, inferior frontal gyrus, and prefrontal cortex. Red reflects increased phase synchrony. Non-significant changes in phase synchrony were set to 0 dB (green). The strongest phase synchronization occurred in the alpha frequency in the posterior parietal, temporal, and anterior cingulate regions after occlusion onset. The posterior parietal cortex also showed gamma phase synchronization with occlusion onset. Strong theta phase synchronization was observed in the right inferior occipital, parietal, and anterior cingulate cortex after occlusion offset. The posterior parietal cortex also showed alpha phase synchronization after the occlusion offset

## Discussion

A likely neural mechanism responsible for the improved dynamic balance performance in Symeonidou and Ferris (2022) is cross modal sensory enhancement from phase resetting in multiple brain areas. Phase resetting increases inter- and intracortical communication and is essential for successfully performing cognitive tasks (Palva et al., 2005; Tass et al., 1998). Stimuli of one modality can cause phase-realignment of other sensory cortices, increasing their processing capacity for subsequent stimuli. In an auditory discrimination task, participants showed an increase in alpha and theta phase synchrony in the auditory cortex when an auditory sound was presented 30-75 ms after a visual stimulus (Thorne et al., 2011). In another study, somatosensory stimulation of the median nerve led to enhanced processing of subsequent auditory stimuli (Lakatos et al., 2007). Most human studies on sensory phase synchrony have focused on audio-visual cross-modal phase resetting or visual-to-auditory phase resetting (Fiebelkorn et al., 2013; Fiebelkorn et al., 2011; Keil & Senkowski, 2018; Lakatos et al., 2009; Mercier et al., 2015; Thorne et al., 2011). However, a study in rats showed that sensory-induced phase resetting can affect the initiation of motor actions (Nicolelis et al., 1995). This suggests that visual stimuli could induce high excitability states in a wide range of brain areas, resulting in increased neural processing of vestibular and sensory inputs, faster responses, and possibly postural adjustments.

Source localization revealed a large number of brain areas with substantive electrocortical activity during beam walking with intermittent visual occlusions. EEG analysis identified electrocortical clusters of independent components in brain areas thought to be directly involved with the control of dynamic balance: occipital (visual processing), temporal (vestibular processing), and sensorimotor cortex (proprioceptive processing) (Horak, 2006; Oliveira et al., 2018; Peterka, 2002; Peterson & Ferris, 2018). There were also clusters in other key brain regions responsible for controlling the body during walking such as the posterior parietal, anterior cingulate, and prefrontal cortex (Gwin et al., 2011; Peterson, Rios, et al., 2018; Sipp et al., 2013). Additional brain regions also appeared in the source localization of EEG clusters, including the inferior frontal gyrus and the posterior cingulate.

Our results implicating both the inferior frontal gyrus and posterior cingulate are in agreement with previous studies implicating those areas in the cortical control of gait (Doi et al., 2017; Sipp et al., 2013). A previous balance study in our lab showed increased theta power in the posterior cingulate when participants lost their balance (Sipp et al., 2013). This region has also been linked to vestibular cortex connectivity (Lopez et al., 2012), high temporal gait variability in healthy individuals, as well as gait abnormalities in patients with Alzheimer’s and Parkinson’s disease (Tian et al., 2017; Wennberg et al., 2017). A Positron Emission Tomography (PET) study comparing imagined locomotion to a rest condition revealed the involvement of the posterior cingulate and the inferior frontal gyrus (Malouin et al., 2003). Another study showed increased posterior cingulate and inferior frontal gyrus activation during gait and dual-tasking (Doi et al., 2017). The inferior frontal gyrus has also been linked to gait speed in single and dual-task conditions and has shown to modulate attentional resources in response inhibition tasks (Hampshire et al., 2010). As the spectral changes in this region were observed after occlusion onset and offset, it is possible that they are related to inhibition of motor responses when visual conditions change.

The prefrontal cortex and anterior cingulate showed fundamentally different patterns of synchronization and desynchronization in relation to the occlusion onset and return of vision compared to the other cortical clusters. A major role for the anterior cingulate is error monitoring (Anguera et al., 2009; Carter et al., 1998), which is important for recognizing deviations from the desired body position (Ahmed & Ashton-Miller, 2004). Studies have shown an increase in anterior cingulate activity when successfully recognizing unstable posture or when postural demands increased in a balance beam walking task (Peterson et al., 2018; Sipp et al., 2013). Specifically, Peterson and Ferris (2018) reported an increase in theta synchronization in the anterior cingulate directly after the onset of a physical perturbation but did not show strong synchronization after a rotation of the visual field. In our results, the anterior cingulate showed a very strong theta synchronization when vision was restored, suggesting that suddenly re-integrating visual sensory information during online balance control more directly involves the anterior cingulate than losing vision. The inter-trial coherence (ITC) data do indicate, however, that the anterior cingulate was involved in the phase resetting that came with both visual occlusion and visual return. In the prefrontal cortex, the gamma desynchronization during occlusion and until ∼500 ms after vision was restored, likely reflects cognitive involvement in body movement. Multiple studies have reported evidence suggesting the prefrontal cortex alters activation levels in response to challenging balance tasks (Basso Moro et al., 2014; Ferrari et al., 2014).

Our EEG evidence suggests that the posterior parietal cortex played a major role in the cortical processing of visual occlusions. Both the event-related spectral perturbation and inter-trial coherence plot show strong responses to the visual occlusion and to the restoration of vision. A recent review of phase synchrony in cross-modal sensory processing (Bauer et al., 2020) discussed the potential role of the primary sensory cortices, superior temporal sulcus, intraparietal sulcus, and prefrontal cortical regions as neural substrates for cross-modal phase synchrony. In locomotion and other whole-body movements, the posterior parietal cortex plays a central role in integrating visual and somatosensory information (Drew & Marigold, 2015; Nordin et al., 2019). In a recent study by Young and colleagues (2020), active transcranial direct-current stimulation (tDCS) of the posterior parietal cortex decreased postural adaptation in an incline-intervention paradigm compared to sham stimulation. The same group found delayed split-belt walking adaptation after inhibiting the posterior parietal cortex contralaterally to the belt with the altered speed (Young et al., 2020b). Both studies highlight the crucial role of the posterior parietal cortex in balance and gait adaptation. In our results, visual occlusion phase synchrony in the posterior parietal cortex extended to the greatest frequency range among the brain regions, including theta, alpha, beta, and gamma frequencies. Gamma neural oscillations have been shown to enhance learning and memory in a wide range of conditions and tasks (Adaikkan & Tsai, 2020). Phase synchrony in the beta and gamma band tend to align action potentials in cortical neurons to synchronize their discharges with good precision (Fries et al., 2007; Gray et al., 1989; Pesaran et al., 2002), suggesting that the posterior parietal cortex may be the driving center of the cross-modal resetting for somatosensory and visual perturbations during locomotion.

We designed our study to investigate how the visual occlusions affected electrocortical dynamics during beam walking, but not to determine how electrocortical processes change with improved balance performance. We found that the onset of the visual occlusion induced increased phase synchrony in multiple electrocortical frequency bands within the sensory (especially the temporal), posterior parietal, and anterior cingulate cortices. However, our study had some limitations which need to be considered when designing future studies. We did not collect enough data from subjects performing unperturbed balance beam walking before and after training to perform independent component analysis on the EEG to test for differences in brain processes between pre- and post-test across participants when they were walking without intermittent visual occlusions. Future research could collect longer trials of pre- and post-test unperturbed balance beam walking to test for potential changes in brain dynamics that occur after the visual occlusions training.

## Conclusion

Our EEG findings suggest that intermittent visual occlusions induce cross-modal phase synchrony in brain areas relevant for sensory processing and balance control. The wide spread phase synchrony could explain enhanced balance training effects found in our previous study (Symeonidou & Ferris, 2022). Future work should examine a larger number of subjects in long-term training and across a range of ages to determine if there are detectable changes in sensory processing electrocortical dynamics when comparing pre- and post-training with intermittent visual occlusions during balance tasks.

## Acknowledgments

We would like to thank Evdokia Ptitsyna and Lourdes Bernandez for their assistance in data collection as well as Roehl Reyes and Crosman Cruz for their assistance in data analysis.

## Author Contributions

E-RS and DF co-designed this study. E-RS acquired and analyzed the data and drafted the manuscript. DF contributed to data interpretation and manuscript drafting. Both authors have read and approved the final manuscript. No one who qualifies for authorship has been omitted. The manuscript has not been published nor accepted for publication elsewhere.

